# Stratifying depression by neuroticism: revisiting a diagnostic tradition using GWAS data

**DOI:** 10.1101/547828

**Authors:** Mark J. Adams, David M. Howard, Michelle Luciano, Toni-Kim Clarke, Gail Davies, W. David Hill, 23andMe Research Team, Major Depressive Disorder Working Group of the Psychiatric Genomics Consortium, Daniel Smith, Ian J. Deary, David J. Porteous, Andrew M. McIntosh

## Abstract

Major depressive disorder and neuroticism share a large genetic basis. We sought to determine whether this shared basis could be decomposed to identify genetic factors that are specific to depression. We analysed two sets of summary statistics from genome-wide association studies of depression (from the Psychiatric Genomics Consortium and 23andMe) and compared them to GWAS of neuroticism (from UK Biobank). First, we used a pairwise GWAS analysis to classify variants as associated with only depression, with only neuroticism, or with both. Second, we estimated partial genetic correlations to test whether the depression’s genetic link with other phenotypes was explained by shared overlap with neuroticism. We found evidence that most genetic variants associated with depression are likely to be shared with neuroticism. The overlapping common genetic variance of depression and neuroticism was negatively genetically correlated with cognitive function and positively genetically correlated with several psychiatric disorders. We found that the genetic contributions unique to depression, and not shared with neuroticism, were correlated with inflammation, cardiovascular disease, and sleep patterns. Our findings demonstrate that, while genetic risk factors for depression are largely shared with neuroticism, there are also non-neuroticism related features of depression that may be useful for further patient or phenotypic stratification.

## Introduction

Major depressive disorder (MDD) is a leading cause of morbidity worldwide, currently affecting approximately 4% of the world’s population^1^. MDD is classified by the World Health Organisation and American Psychiatric Association according to its severity, its recurrence or chronicity, and the presence or absence of psychotic symptoms. This approach aims to maximise the reliability of MDD’s diagnosis while being agnostic about its underlying aetiology until robust evidence of causal mechanisms can be used to stratify the condition.

Historically, depression had been classified on the basis of pre-existing emotional instability into ‘neurotic’ and ‘endogenous’ forms^2^. ‘Neurotic depression’ was diagnosed in the presence of pre-existing emotional instability and was reported to occur in younger individuals who more frequently expressed suicidal ideation. ‘Endogenous depression’, in contrast, was characterised by more frequent melancholic symptoms, including disrupted sleep, impaired appetite, diurnal variation of mood and impaired cognition. Individuals with endogenous depression were described as more responsive to antidepressant treatments^3^.

The nosological status of neurotic and endogenous depression has, however, been very controversial^4^. Some studies have found empirical support for the neurotic/endogenous division based upon self-report and clinical data^5^, whereas other studies found little empirical support for the distinction and noted that individuals with low mood responded to antidepressant treatments regardless of their subtype^6^.

The tendency toward emotional instability, or neuroticism, is a robustly and consistently replicated dimension of personality that is relatively stable over time^7^. Neuroticism features in the most widely accepted theory of personality structure, the Five Factor model, alongside openness to experience, extraversion, conscientiousness and agreeableness^8^. Trait neuroticism (sometimes also labelled as emotionality or its opposite, emotional stability) is identified consistently and is composed of items reflecting low mood, stress sensitivity, irritability, and emotional control^9^.

Twin, family, and genomic studies have shown that population variation in neuroticism and liability to depression are conferred by both genetic and environmental risk factors^10–15^. While earlier studies^16^ had difficulty discovering specific genetic variants for depression, possibly partly because of heterogeneity^17^, recent studies have identified genome-wide significantly associated loci either by focussing on more rigorously defined MDD phenotypes^18^ or by meta-analyzing case-control samples with larger sample-size studies that use broader depression phenotypes^19–22^. Recent studies of neuroticism have also identified over one hundred genome-wide significant loci^23,24^. Confirming earlier findings from biometric studies^25,26^, these studies have shown that depression and neuroticism have a genetic correlation of approximately 0.7 and thus share a significant proportion of their genetic architecture^23^. Since this genetic correlation is sizeable but still less than 1, it suggests that depression and neuroticism are either both unreliable measures of the same underlying genetic predisposition or it indicates that depression and neuroticism may not be identical in terms of their genetic architecture. This may be because the effect sizes of each variant^27^ differ between depression and neuroticism, which includes the possibility that some variants affect one trait but not the other. A third (non-exclusive) possibility is that neuroticism or one of its facets is directly related to only a subtype of depression and that other subtypes will have distinct genetic aetiologies.^24,28^

In the current study we sought to compare genome-wide association summary statistics for depression and neuroticism to test whether there are specific genetic contributions to depression that are independent of neuroticism. First, we sought to identify loci that conferred a higher risk of depression, but not higher neuroticism, and vice versa. We then annotated the function of any associated loci. Secondly, we sought to determine whether depression and neuroticism have any unique genetic correlations with other traits and disorders after adjustment for the other variable.

## Methods

### Data sources

We used depression summary statistics from the Psychiatric Genomics Consortium (PGC) ^20^ and from the 23andMe cohort^19^. We obtained PGC MDD summary statistics that excluded the UK Biobank cohort. The version of the meta-analysis that we used was conducted on 116,404 cases and 314,990 controls (N= 431,394, prevalence = 0.27) and included 9,030,847 variants. We compared the depression GWASs to summary statistics from a GWAS of neuroticism in UK Biobank (N = 329,821) with 18,485,883 variants.^23^

### Pairwise GWAS of depression and Neuroticism

In order to identify loci that contribute to variation in liability to depression, but not neuroticism, we used pairwise GWAS^29^ to jointly analyse summary statistics from depression and neuroticism. We used depression summary statistics from the Psychiatric Genomic Consortium MDD2 meta analysis^20^ that excluded summary statistics from the UK Biobank sample (N cases = 116,404; N controls = 314,990); for neuroticism we used previously published summary statistics from the UK Biobank sample^23^. We used the munge_sumstats.py tool^30^ to convert the summary statistics to z-scores and to align and filter effect alleles to a common reference (HapMap 3 SNPs). Using these filtered summary statistics, the *gwas-pw* program^29^ models the probability that each locus is associated with only one of the two traits, the probability that the locus has a shared association with both traits, and the probability that each of the genomic regions contains separate loci that are associated with each trait. We analysed the summary statistics by splitting them into genomic segments (1702 in total) that were approximately independent based on linkage disequilibrium.^31^ Although we used summary statistics from different studies, to correct for potential cohort overlap we estimated the expected correlation of effect sizes if there were no loci associated with both traits. To do this we estimated the probability that each genomic segment contained a locus that was significantly associated with at least one of the traits, using the fgwas program.^32^ We then retained segments that had a posterior probability of association < 0.2 with either trait. Using these non-associated segments, we calculated the correlation of effect sizes across the depression and neuroticism summary statistics, and then supplied this correlation coefficient to gwas-pw. The gwas-pw program performs a Bayesian test on each genomic region to estimate the probability of unique or overlapping association signals from the two sets of GWAS summary statistics^29^.

Using the *gwas-pw* output, we categorized each segment as being associated with depression only, neuroticism only, both traits, or neither trait. We defined depression-only segments as those that had the highest posterior probability associated with only depression and that also had a genome-wide significant hit in the original GWAS but did not contain a genome-wide significant hit for neuroticism (and vice versa for neuroticism-only segments). We defined segments that were associated with neither trait as those that had a total posterior probability of association < 0.2 and that did not contain any genome-wide hits for either traits. To examine the depression-only signal, we excluded segments associated with neuroticism or with both traits from the depression summary statistics, clumped SNPs that were in LD (r^2^ > 0.1) and within 3Mb, then used MAGMA ^33^ to identify significantly associated (p < 2.77 × 10^-6^) genes and conducted GWAS catalog lookups using FUMA ^34^. We then conducted the same set of analyses on the neuroticism-only segments from the neuroticism summary statistics and the segments associated with both depression and neuroticism from the depression summary statistics.

### Genetic correlations with depression adjusted for neuroticism

We used cross-trait genetic correlations to identify traits that were related to depression after removing their shared genetic effects with neuroticism. To start, we first obtained estimates of genetic correlations from GWAS summary statistics that were downloaded from LD Hub^35^. The LD Hub results contained information on genetic correlations with MDD from the Psychiatric Genomics Consortium^16^ and neuroticism from the Genetics of Personality Consortium^36^. We selected 23 traits of interest that were nominally genetically correlated (p < 0.01) with either MDD or neuroticism. These included psychiatric disorders, personality traits, cardiovascular and inflammatory diseases, anthropometric and life-history traits, education, and lifestyle factors. We supplemented this list of suggestive traits with body mass index since obesity has been identified as a potential stratifying factor for depression^37^. For each trait of interest, we calculated its genetic correlation using LD Score regression ^30^ with an independent set of GWAS summary statistics for MDD and Neuroticism. For depression we used data from 23andMe^19^ and for Neuroticism we used data from UK Biobank^38^.

Our goal was to determine whether depression’s genetic correlation with each trait of interest is (1) explained by the genetic architecture shared between depression and neuroticism or (2) specific to depression and independent of neuroticism. We aimed to test between these two possibilities by taking each trait’s genetic correlation with depression and partitioning out shared genetic overlap with neuroticism to estimate a partial genetic correlation. To make statistical inferences about the partial genetic correlations, we needed to account for the joint uncertainty of the LD score genetic correlation estimates among depression, neuroticism, and the other traits and to smooth the genetic correlation matrix to ensure it was invertible. To do this, we defined a model that treated each pairwise genetic correlation estimate as a sample from a true underlying genetic correlation matrix between all traits:

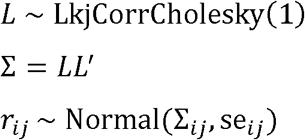

where ∑ is the true correlation matrix; *L* is drawn from a LKJ distribution^39^ with a shape parameter η = 1 (which yields a prior distribution that is uniform for all correlations between −1 and 1); *r*_*ij*_ is the LD score genetic correlation between traits *i* and *j* and se_ij_ is its standard error. We implemented the model in the Bayesian modelling language Stan^40^ and ran it using a ‘No-U-Turn sampler’ with 4 chains for 2000 iterations and a warmup of 1000 iterations. We calculated the partial genetic correlation between depression and trait *i* that removed the shared overlap with neuroticism as:

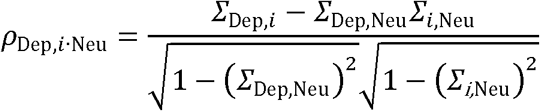

To determine if neuroticism explained all of the genetic correlation of each trait with depression, we tested whether the partial genetic correlation was significantly different from zero. To analyze the specificity of the genetic correlation with respect to depression and the attenuation of the partial genetic correlation, we tested the magnitude of the partial correlation adjustment, *∑*_Dep,Neu_*∑*_*i*,Neu_ ≠ 0. We repeated the same procedure to estimate the partial genetic correlation between each trait and neuroticism after removing shared overlap with MDD.

## Results

### Pairwise GWAS of depression and Neuroticism

We used pairwise GWAS^29^ between depression and neuroticism to partition genomic segments with association signals for MDD only, for neuroticism only, for both traits, or that contained different associations for each trait. We used summary statistics from the PGC for depression^20^ and from UK Biobank for neuroticism^23^. The correlation in beta coefficients from the summary statistics in genomic segments that were not associated with either trait was *r* = 0.005, suggesting there was no undetected sample overlap between the two studies. We identified 9 genomic segments containing loci that influence depression but do not associate with neuroticism (Figure 1, Table 1). This represented 24% of the total 37 genomic segments that contained loci associated with depression. The analysis indicated that there were three genomic segments that contained separate associations for depression and neuroticism (Supplementary Table S3 and Figures S2–4). There were 25 GWAS hits for depression that were also associated with neuroticism; in addition, there were 20 more regions that, while below the threshold of genome-wide significant for depression, the pairwise analysis indicated were associated with both depression and neuroticism. Finally, there were also 40 genomic segments that were significantly associated only with neuroticism (Supplementary Table S1).

**Table 1:**
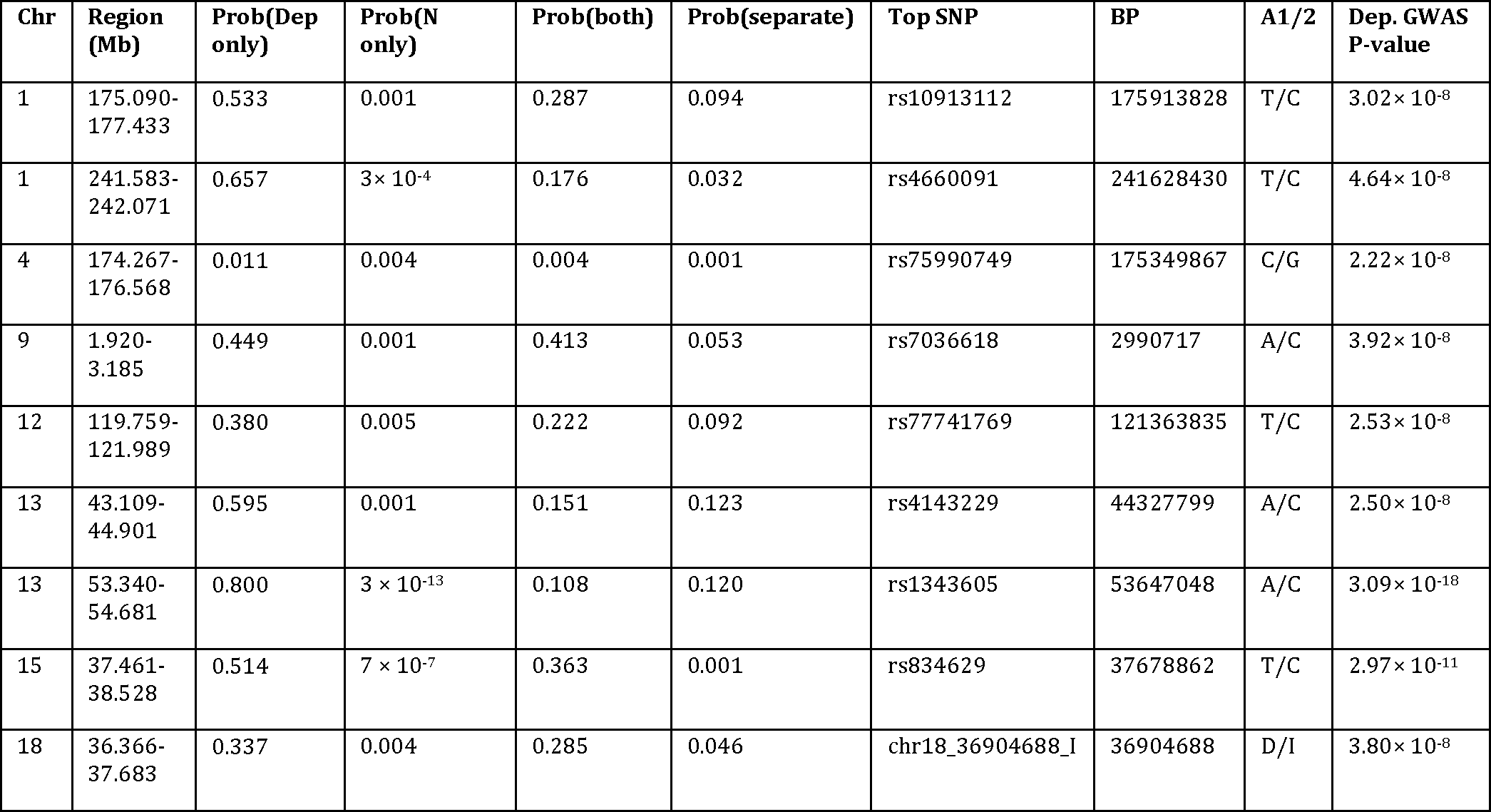
Genomic regions that are likely associated with depression but not with neuroticism that also contain a genome-wide significant SNP for depression. For each segment, the posterior probability of association (higher value means more probable) with depression only [Prob(Dep only)] is compared with Neuroticism only [Prob(N only)], with the same association signal for both traits (Prob(both)), or with separate association signals effecting each trait (Prob(separate)). For each region, the top SNP associated with MDD is listed along with its basepair position (BP) and GWAS *p*-value.

| Chr | Region (Mb) | Prob(Dep only) | Prob(N only) | Prob(both) | Prob(separate) | Top SNP | BP | A1/2 | Dep. GWAS P-value |
| --- | --- | --- | --- | --- | --- | --- | --- | --- | --- |
| 1 | 175.090-177.433 | 0.533 | 0.001 | 0.287 | 0.094 | rs10913112 | 175913828 | T/C | $3.02 \times 10^{-8}$ |
| 1 | 241.583-242.071 | 0.657 | $3 \times 10^{-4}$ | 0.176 | 0.032 | rs4660091 | 241628430 | T/C | $4.64 \times 10^{-8}$ |
| 4 | 174.267-176.568 | 0.011 | 0.004 | 0.004 | 0.001 | rs75990749 | 175349867 | C/G | $2.22 \times 10^{-8}$ |
| 9 | 1.920-3.185 | 0.449 | 0.001 | 0.413 | 0.053 | rs7036618 | 2990717 | A/C | $3.92 \times 10^{-8}$ |
| 12 | 119.759-121.989 | 0.380 | 0.005 | 0.222 | 0.092 | rs77741769 | 121363835 | T/C | $2.53 \times 10^{-8}$ |
| 13 | 43.109-44.901 | 0.595 | 0.001 | 0.151 | 0.123 | rs4143229 | 44327799 | A/C | $2.50 \times 10^{-8}$ |
| 13 | 53.340-54.681 | 0.800 | $3 \times 10^{-13}$ | 0.108 | 0.120 | rs1343605 | 53647048 | A/C | $3.09 \times 10^{-18}$ |
| 15 | 37.461-38.528 | 0.514 | $7 \times 10^{-7}$ | 0.363 | 0.001 | rs834629 | 37678862 | T/C | $2.97 \times 10^{-11}$ |
| 18 | 36.366-37.683 | 0.337 | 0.004 | 0.285 | 0.046 | chr18_36904688_I | 36904688 | D/I | $3.80 \times 10^{-8}$ |

**Figure 1.**
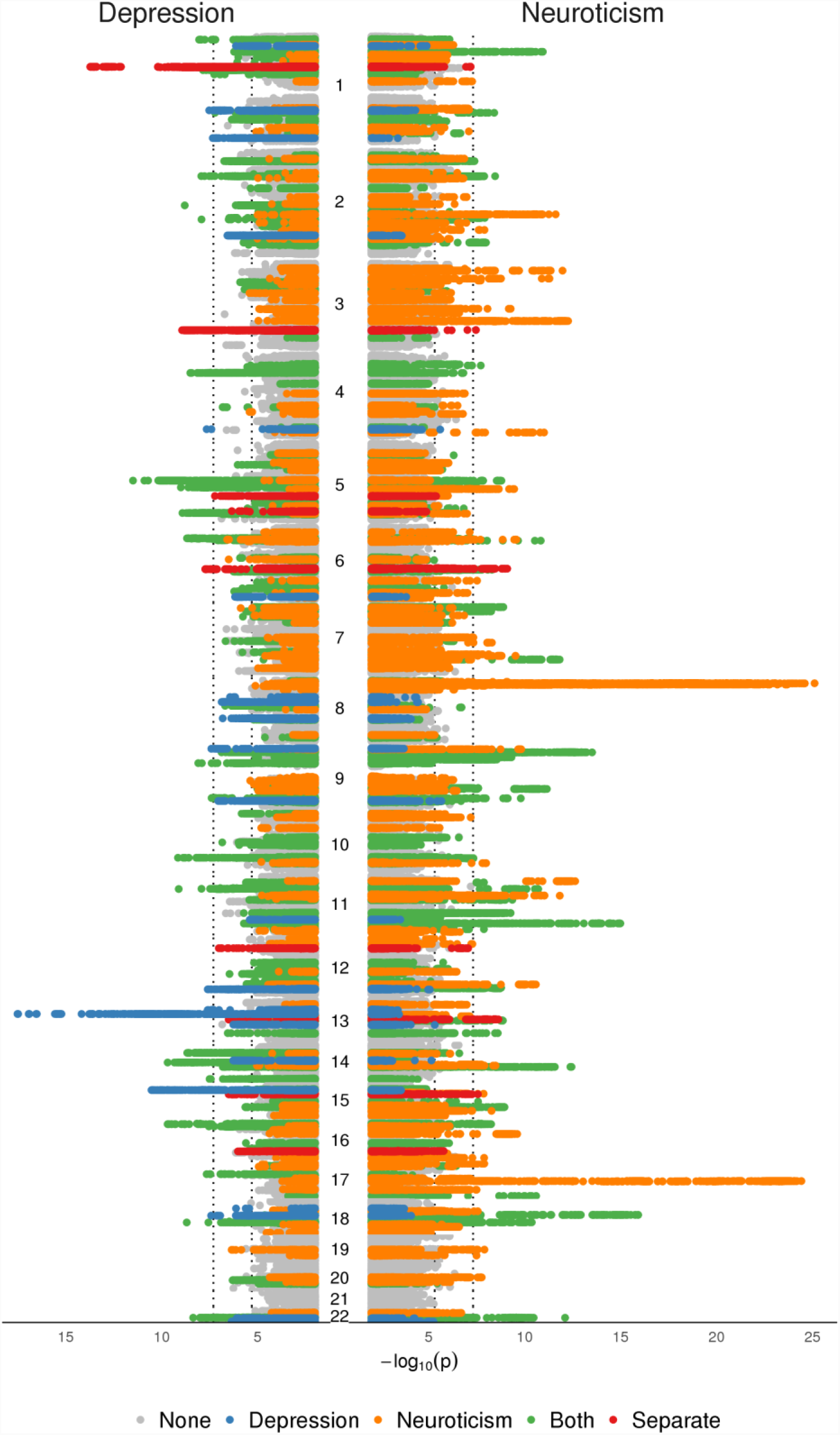
Manhattan plot of pairwise GWAS of depression and neuroticism with genomic segments partition by association with depression-only (blue), neuroticism only (orange), both traits (green), separate associations (red), or neither trait (gray).

We used MAGMA^33^ to identify genes associated (*p* < 2.77 × 10^-6^) with the partitioned genomic segments. There were 30 genes significantly associated with depression only (Supplementary Table S2), 203 genes associated with neuroticism only (Supplementary Table S3), and 104 genes associated with both depression and neuroticism (Supplementary Table S4). We used FUMA ^34^ to conduct GWAS catalogue lookups on these gene sets (Supplementary Table S6). MDD was uniquely associated with gene sets that are linked to N-glycan levels (*p* = 1.1 × 10^-8^) and to coronary heart disease (*p* = 3.2 × 10^-3^). Neuroticism-only gene sets were related to traits such as intracranial volume, Parkinson’s disease, and HDL cholesterol. The gene sets shared by depression and neuroticism were related to cross-disorder psychiatric traits, coffee consumption, and epilepsy, among other traits (Supplementary S5).

### Partial genetic correlations with depression removing Neuroticism

We used LD Score regression results from LD Hub^35^ to identify traits that were significantly genetically correlated with depression or neuroticism and then tested whether genetic overlap with neuroticism explained each trait’s genetic correlation with depression (and vice versa) using an independent set of summary statistics for depression (23andMe) and neuroticism (UK Biobank). The LD score genetic correlation between 23andMe depression and UKB neuroticism was 0.602±0.025 SE (*p* = 1.65 × 10^- 125^). Figure 2 shows the unadjusted genetic correlation of each trait of interest with depression and with neuroticism against the partial genetic correlation with depression where the shared overlap with neuroticism has been removed (Depression**·**Neuroticism) and with neuroticism where overlap with depression has been removed (Neuroticism**·** Depression).

**Figure 2.**
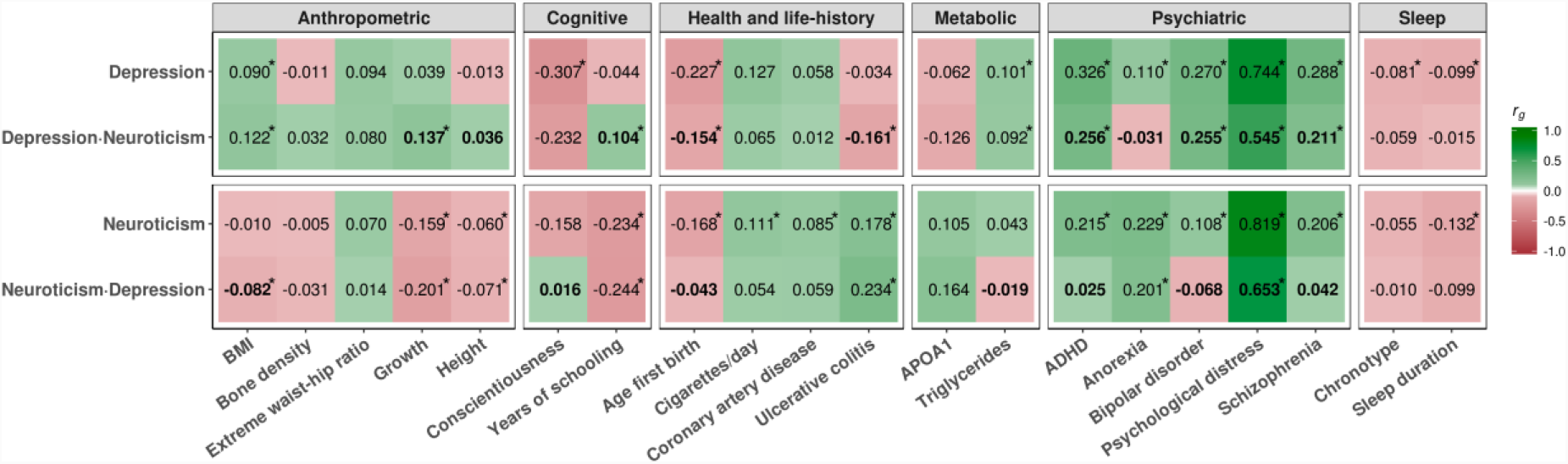
Genetic correlations of traits with depression and with neuroticism and partial genetic correlations with MDD adjusting for neuroticism (Depression**·**Neuroticism) and with neuroticism adjusting for MDD (Depression **·**Neuroticism). Estimates are marked with ‘*’ if the genetic correlation or partial genetic correlation is significantly different from 0 (adjusted for multiple testing). Each partial genetic correlation is in bold if it is significantly different from the full genetic correlation.

We tested whether each partial genetic correlation was different from 0 and whether it was attenuated from its unadjusted value. Removing genetic overlap with neuroticism completely attenuated the genetic correlation of MDD with anorexia. Removing genetic overlap with neuroticism substantially attenuated the genetic correlation of MDD with attention deficit hyperactivity disorder, psychological distress, schizophrenia, and age of first birth. Depression’s genetic correlations with bipolar disorder, body mass index, conscientiousness, chronotype (“morningness”), and triglycerides were not attenuated by the removal of shared genetic overlap with neuroticism. For neuroticism, genetic overlap with depression completely explained its genetic correlation with ADHD, bipolar disorder, schizophrenia, and conscientiousness. Neuroticism’s genetic correlation with psychological distress was also substantially, but not completely, attenuated after factoring out shared overlap with depression while neuroticism’s genetic correlations with pubertal growth, height, years of schooling, and ulcerative colitis were unchanged. For several traits the partial genetic correlation was significantly different from zero even though the unadjusted genetic correlation with depression or neuroticism was not; this occurred when the trait had a sufficiently stronger genetic correlation with, e.g., neuroticism (in the case of years of schooling) or when the its genetic correlation with neuroticism went in the opposite direction (ulcerative colitis, pubertal growth).

## Discussion

We conducted complementary analyses to separate the specific genetic features of depression from those that overlap with neuroticism. The pairwise GWAS analysis using depression results from the PGC and neuroticism results from UK Biobank revealed 9 genomic regions that were significantly associated with depression but not with neuroticism. Several of the associated regions contained genes of known function: rs10913112 is downstream of *RFWD2* (ring finger and WD repeat domain 2), a gene that can promote tumor growth^41^; rs4660091 is near the fumarate hydratase gene (*FH*) which is involved in the Krebs cycle; rs4143229 is in an intron of the ecto-NOX disulfide-thiol exchanger 1 gene (*ENOX1*) which is expressed in the nervous systems and has been implicated in autoimmune disorders^42^; and rs1343605 is near *OLFM4*, a gene that has been linked to depression^20^, inflammation, cancer^43^.

The pairwise GWAS also suggested three regions that contained separate association signals for depression and neuroticism. For example, a region (71.6-74.3Mb on chromosome 1, Figure S1) contained two association signals for depression. The first signal (rs1460942) was shared with neuroticism and was close to the neuronal growth regulator 1 (*NEGR1*) gene. A second association signal (rs12129573) was in the intron of an uncharacterized non-protein coding RNA (*LOC105378800*). In a further region, a signal was found (97.8-100.6Mb on chromosome 6) that showed a clear separation between the association signals for each trait (Figure S3). The SNP associated with depression (rs12202410) was near the F-box and leucine rich repeat protein 4 (*FBXL4*) gene which is related to energy homeostasis ^44^.

Using a GWAS catalog lookup with FUMA^34^ on the pairwise GWAS results, we found that genes associated with depression, but not with neuroticism, were also associated with glycosylation and coronary heart disease. This suggests that there may be subtypes of depression involving inflammation and cardiovascular disease that are separate from subtypes of depression associated with neuroticism. In contrast, genes that were associated with both depression and neuroticism were also associated with psychological, behavioral, and psychiatric traits such as schizophrenia, autism spectrum disorder, and intelligence, suggesting that they may influence behaviour and cognitive function more generally.

Differential association with other traits, once shared overlap with neuroticism was accounted for, was also shown in our analysis of LD score genetic correlations. We modelled the underlying genetic correlation matrix from pairwise LD score estimates, which is similar to the stage 1 estimation step in Genomic SEM^45^ or to calculating overlap in genetic correlations using GWIS^46^. Our model is particularly useful when the parameter of interest (such partial correlation coefficients, as in our study) can be calculated directly from the estimated genetic matrix rather than via a more complex estimation procedure. In our analysis of the genetic correlation matrix, we found that depression (unlike neuroticism) was uniquely correlated with bipolar disorder, body mass index, chronotype, and triglyceride levels. The genetic architecture of depression that is shared with neuroticism explained all of most of the genetic correlation of depression with ADHD, anorexia, psychological distress, and schizophrenia, among other traits.

The major limitation of our study is that by using summary statistics from genome-wide association studies, we were only able to assess the overlap in genetic architecture between depression and neuroticism that arises from common variants. Even biobank-sized samples of millions of participants can be underpowered for detecting associations with rare variants unless such variants have very large effect sizes {Visscher, 2017 #2049}.

Neuroticism is a major risk factor for depression^47,48^ and the two traits are strongly genetically correlated^23,25,49^. Our results confirm that the majority of specific genetic variants associated with depression are shared with neuroticism. However, we also identified several association signals and overlap with other traits that were unique to depression. In particular, the pairwise GWAS analysis both identified new associations with depression and separated out associations that were specific to depression. This entails that that depression and neuroticism are not *just* noisy measures of the same underlying liability. If these differences represent distinct genetic subtypes of depression, then most cases of depression will stem from this neuroticism-depression nexus, while a smaller proportion may have aetiologies that are distinct from neuroticism. Some of these associations, such as between depression, chronotype, and metabolic phenotypes, are suggestive of endogenous depression’s features but do not point to all the characteristics of this previously described subtype. Neither depression^28^ nor neuroticism^24^ is completely genetically homogeneous. Because the common variant genetic overlap is high between different forms of assessing depression (clinically ascertained, brief questionnaire, hospital records)^20,50^, our results suggest that much of the shared genetic variance, where depression and non-depression are on a continuum, is wrapped up in the neuroticism-depression nexus, and that studies based on a more refined, symptom-level analysis may be more revealing for non-neuroticism related forms of depression.

### Data availability

The full MDD summary statistics from Hyde et al. are made available through 23andMe to qualified researchers under an agreement with 23andMe that protects the privacy of the 23andMe participants. Please visit https://research.23andme.com/collaborate for more information and to apply to access the data. Summary statistics from the Psychiatric Genomics Consortium are available for download from http://www.med.unc.edu/pgc/. Summary statistics for Neuroticism in UK Biobank are available from the Centre for Cognitive Aging and Cognitive Epidemiology http://www.ccace.ed.ac.uk/node/335.

## Acknowledgements

We thank the following members of the 23andMe Research Team: Michelle Agee, Babak Alipanahi, Adam Auton, Robert K. Bell, Katarzyna Bryc, Sarah L. Elson, Pierre Fontanillas, Nicholas A. Furlotte, David A. Hinds, Karen E. Huber, Aaron Kleinman, Nadia K. Litterman, Jennifer C. McCreight, Matthew H. McIntyre, Joanna L. Mountain, Elizabeth S. Noblin, Carrie A.M. Northover, Steven J. Pitts, J. Fah Sathirapongsasuti, Olga V. Sazonova, Janie F. Shelton, Suyash Shringarpure, Chao Tian, Joyce Y. Tung, Vladimir Vacic, and Catherine H. Wilson.

We acknowledge support from the Wellcome Trust Strategic Award “STratifying Resilience and Depression Longitudinally” (STRADL) (Reference 104036/Z/14/Z) and the MRC Mental Health Data Pathfinder Award (Reference MC_PC_17209). IJD is supported by the Centre for Cognitive Ageing and Cognitive Epidemiology, which is funded by the Medical Research Council and the Biotechnology and Biological Sciences Research Council (MR/K026992/1). The PGC has received major funding from the US National Institute of Mental Health and the US National Institute of Drug Abuse (U01 MH109528 and U01 MH1095320).

IJD is a participant in UK Biobank. Members of the 23andMe Research Team are employees of 23andMe, Inc. The other authors declare they no other conflicts of interest.

This work has made use of the resources provided by the Edinburgh Compute and Data Facility (ECDF) (http://www.ecdf.ed.ac.uk/).

## Supplementary Information

### Target traits used in partial genetic correlation analysis

Traits were identified as nominally genetically correlated with either major depressive disorder or neuroticism based on results in LD Hub^1^. Summary statistics from GWAS on the following traits were used to estimate genetic correlations: ADHD^2^, Anorexia Nervosa^3^, Bipolar disorder^4^, Psychological distress^5^, Schizophrenia^6^, Conscientiousness^7^, Coronary artery disease^8^, Ulcerative colitis^9^, Age of first birth^10^, Body mass index^11^, Extreme waist-to-hip ratio^12^, Femoral neck bone mineral dens.^13^, Height^14^, Pubertal growth^15^, Chronotype^16^, Sleep duration^16^, Apolipoprotein A-I^17^, Triglycerides^18^, Cigarettes smoked per day^19^, Years of schooling^20^.

**Figures S1–S3:** Manhattan plots for regions identified by pairwise GWAS analysis to have separate association signals for MDD (plotted as −log_10_(p)) and Neuroticism (plotted as log_10_(p)). Color of each SNP represents linkage (R2) with the top SNP from each GWAS in this region.

**Figure S1.**
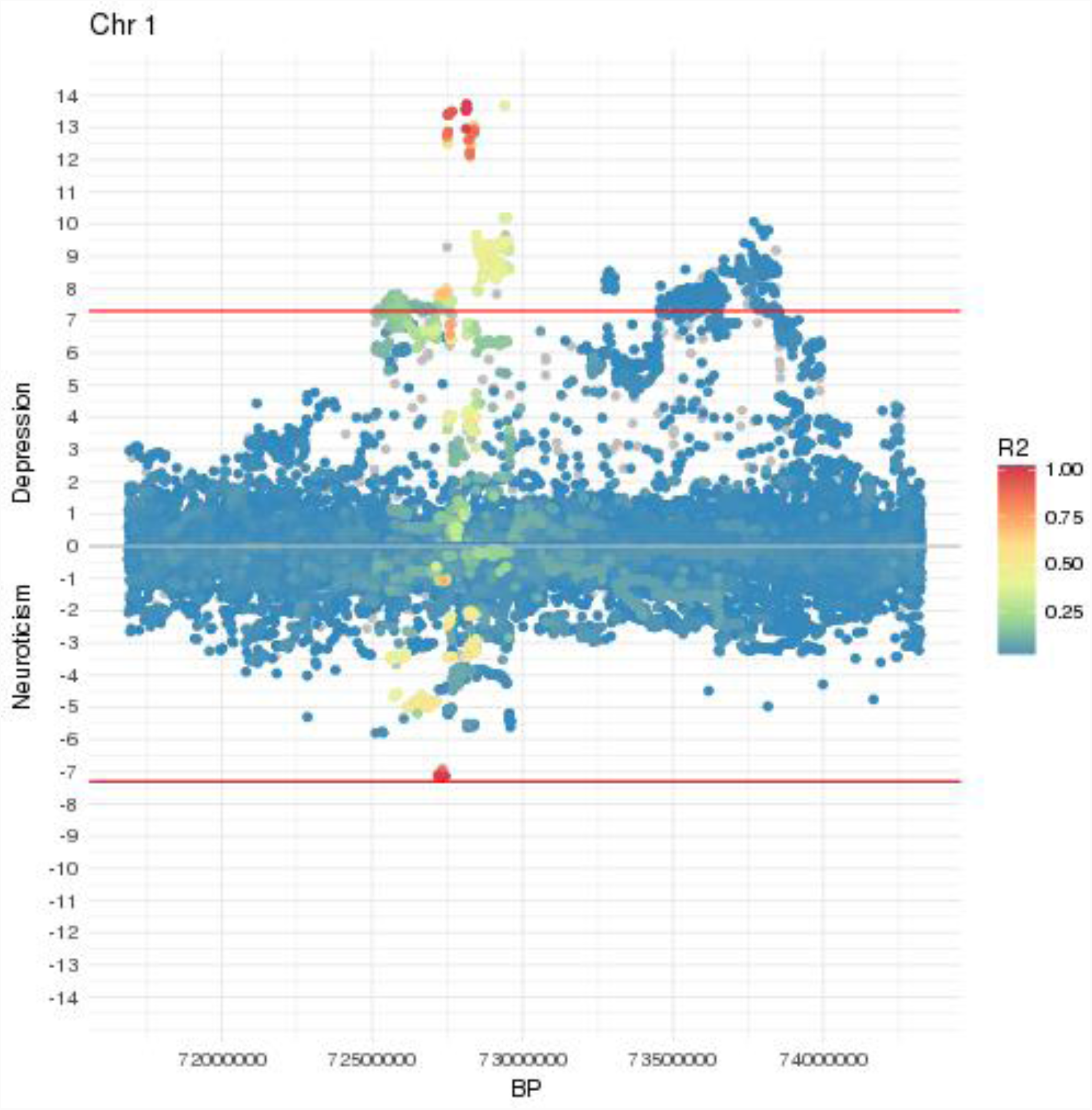
Region 71.6-74.3Mb on chromosome 1

**Figure S2.**
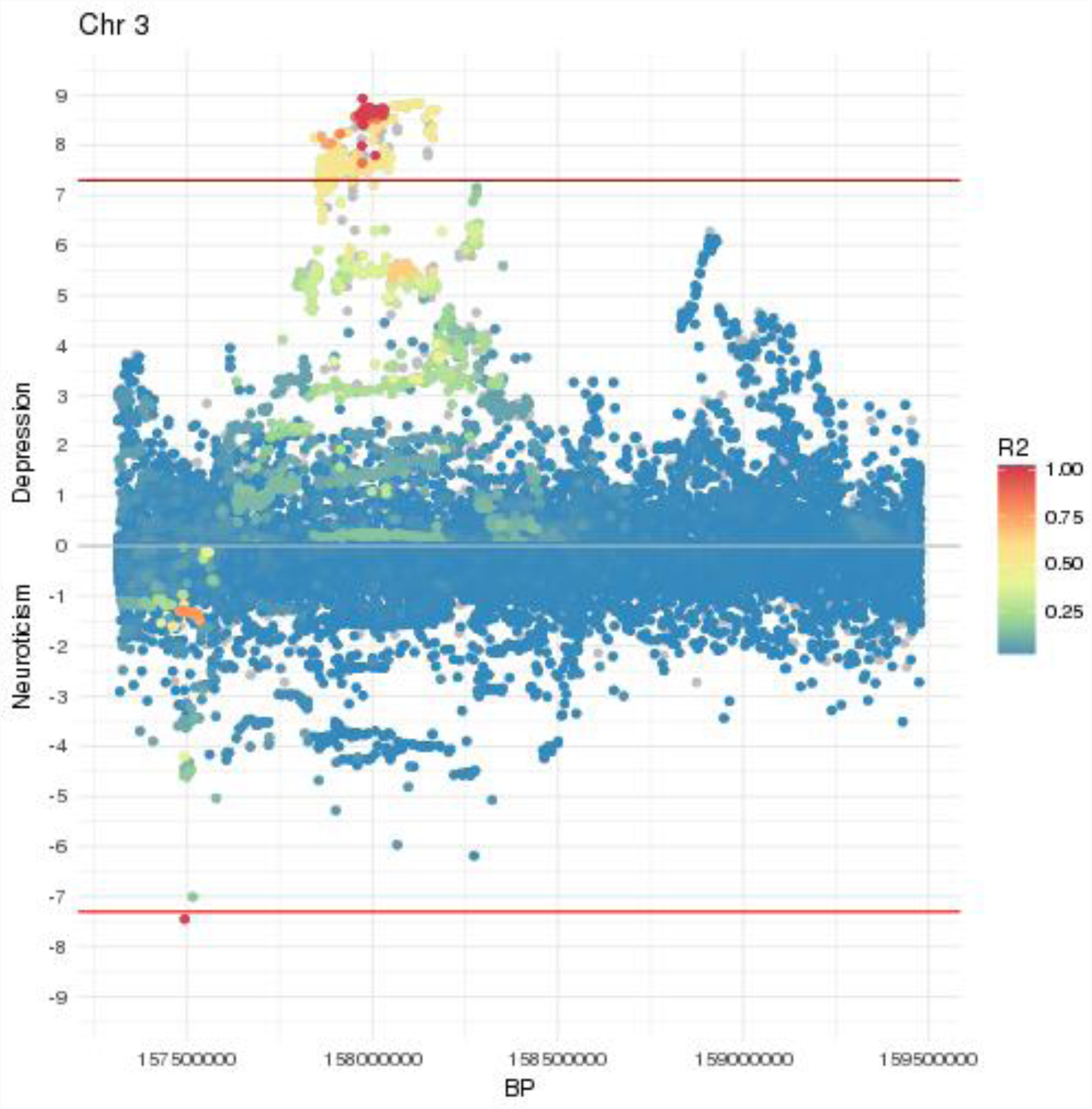
Region 157.3-159.5MB on chromosome 3.

**Figure S3.**
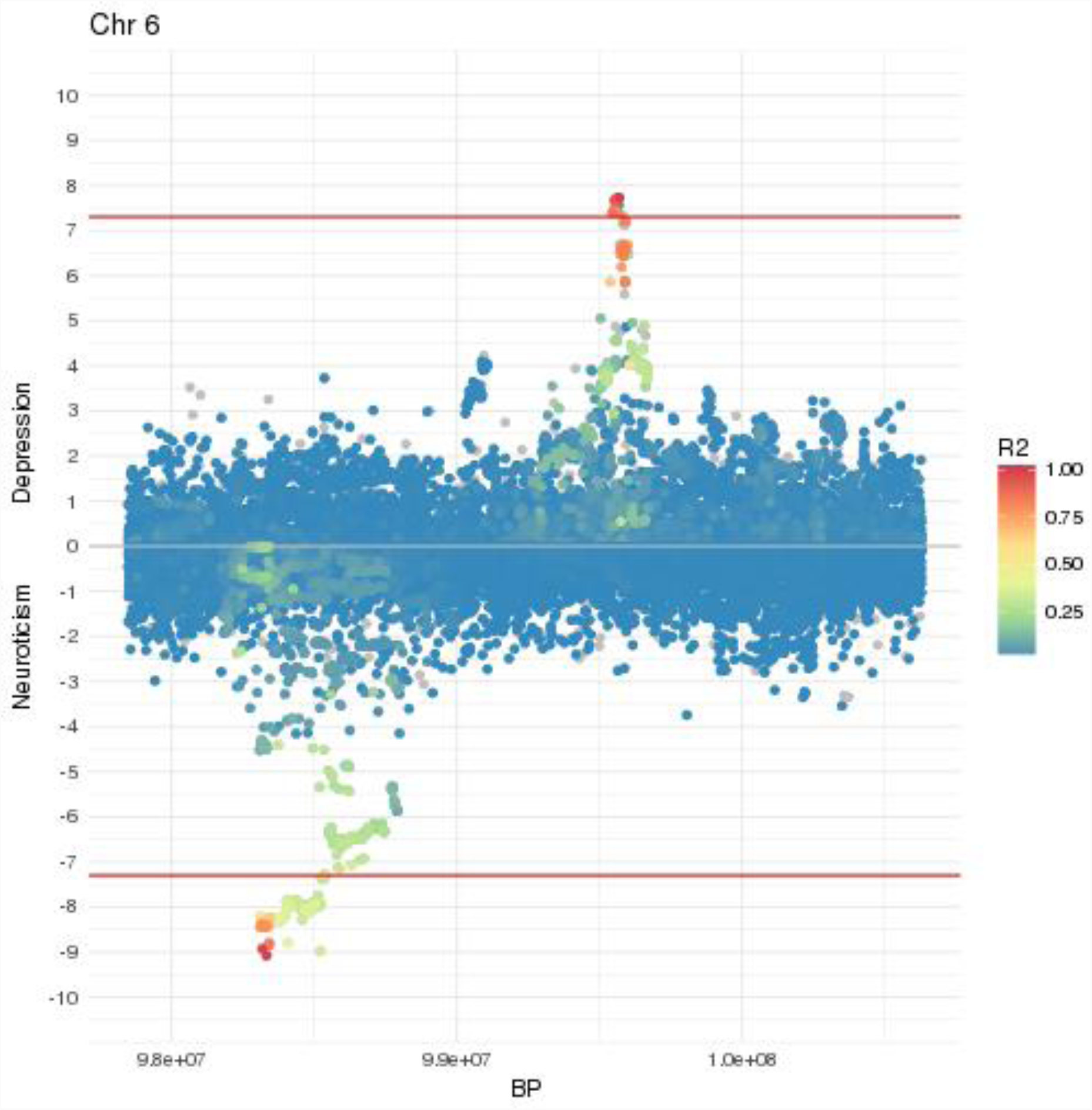
Region 97.8-100.6Mb on chromosome 6

